# A Ca^2+^-regulated deAMPylation switch in human and bacterial FIC proteins

**DOI:** 10.1101/323253

**Authors:** Simon Veyron, Giulia Oliva, Monica Rolando, Carmen Buchrieser, Gérald Peyroche, Jacqueline Cherfils

## Abstract

FIC proteins regulate molecular processes from bacteria to humans by catalyzing post-translational modifications (PTM), the most frequent being the addition of AMP using ATP as a cofactor, a reaction coined AMPylation. In a large group of AMPylating FIC proteins, which includes single-domain bacterial FIC proteins and animal FICD/HYPE, AMPylation is intrinsically repressed by a structurally conserved glutamate. Curiously, FICD departs from previously characterized bacterial FIC proteins in that it acts bifunctionally to AMPylate and deAMPylate its target, the endoplasmic reticulum BiP/GRP78 chaperone. BiP is a key component of the unfolded protein response (UPR), and is AMPylated under normal conditions where its activity is low, while its activation correlates with its deAMPylation. Currently, a direct signal regulating AMPylation efficiency in bacterial and animal FIC proteins has not been identified. Here, we addressed this question for a FIC protein from the bacterial pathogen *Enterococcus faecalis* (EfFIC) and for human FICD. We discover that EfFIC catalyzes both AMPylation and deAMPylation within the same active site, suggesting that the conserved glutamate is the signature of AMPylation/deAMPylation bifunctionality. Crystal structures and PTM assays identify a multi-position metal switch implemented by the glutamate, whereby EfFIC uses Mg^2+^ and Ca^2+^ to control AMPylation and deAMPylation differentially without conformational change. Remarkably, we find that variations in Ca^2+^ levels also tune deAMPylation of BiP by human FICD. Together, our results identify metals as diffusible signals that can regulate bifunctional FIC proteins directly, and they suggest that FICD has features of an enzymatic sensor of Ca^2+^ depletion, a hallmark of the UPR.

## Introduction

In less than a decade, FIC proteins have emerged as a large family of enzymes controling the activity of target proteins by post-translationally modifying them with phosphate-containing compounds (reviewed in ^1,2, 3, 4^). These proteins are characterized by the presence of a conserved FIC domain, which carries out the post-translational modification (PTM) of a Tyr, Ser or Thr residue in a target protein ^5, 6, 7, 8, 9, 10, 11^. The most frequent PTM reaction catalyzed by FIC enzymes is the addition of AMP using ATP as a cofactor, coined AMPylation or adenylylation. This PTM activity was originally discovered in toxins from bacterial intracellular pathogens ^12^ and was later identified in the toxin component of bacterial toxin/antitoxins modules (e.g. Bartonella VbhT/VbhA, ^7^) and in various other bacterial FIC proteins of unknown functions, including single-domain (e.g. Neisseria FIC ^7^) and larger (e.g. Clostridium FIC ^11^) FIC proteins. Metazoans possess a single FIC protein, FICD/HYPE (FICD hereafter), which controls the reversible AMPylation of BiP/GRP78, an Hsp70 chaperone localized in the endoplasmic reticulum (ER) ^13, 14, 15^. BiP is a major component of the unfolded protein response (UPR), a major pathway whereby cells respond to ER stress (reviewed in ^16^). AMPylation of BiP reduces its affinity for unfolded protein clients ^17^ and inversely correlates with the burden of unfolded proteins ^15^. In a recent twist, FICD was shown to act bifunctionally in both AMPylation and deAMPylation of BiP and to carry out deAMPylation as its primary enzymatic activity in vitro ^18^.

All PTM reactions catalyzed by FIC proteins use a motif of conserved sequence for catalysis, the FIC motif, which carries an invariant histidine that is critical for nucleophilic attack of the cofactor by the target residue, and an acidic residue (aspartate or glutamate) that binds an Mg^2+^ ion to stabilize the negative charges of the cofactor phosphates at the transition state (reviewed in ^1,2, 3, 4^). A large group of FIC proteins also features a structurally conserved glutamate that represses AMPylation by protruding into the catalytic site from either N-terminal elements, as in human FICD ^10^, *Clostridum difficile* FIC ^11^ or *Shewanella oneidensis* FIC ^19^, or from a C-terminal α-helix as in single-domain FIC proteins from *Neisseria meningitidis* ^7^ and *Helicobacter pylori* (PDB 2F6S). Crystal structures showed that this glutamate prevents ATP from binding in a catalytically-competent conformation ^7,11^, while its mutation into glycine creates space for the γ-phosphate of ATP ^7^ and has been consistently shown to increase AMPylation activities *in vitro* and in cells (reviewed in ^20^). These observations led to propose that this conserved glutamate represses AMPylation by impairing the utilization of ATP as a donor for AMP, hence that it must be displaced to allow productive binding of ATP ^7^. In *N. meningitidis* FIC (NmFIC), upregulation of the AMPylation activity was proposed to occur upon changes in the toxin concentration ^21^. In this scheme, NmFIC is in an inactive tetrameric state at high concentration, which is further stabilized by ATP, while its dilution promotes its conversion into a monomeric state, leading to activation by displacement of the inhibitory glutamate followed by autoAMPylations that reinforce AMPylation efficiency ^21^. Whether a similar mechanism applies to other glutamate-bearing FIC proteins has not been investigated. Remarkably, in bifunctional human FICD, the autoinhibitory glutamate is critical for deAMPYlation of the BiP chaperone ^18^, a reaction whereby FICD contributes to the activation of BiP under ER stress conditions. Currently, a signal that represses the deAMPylation activity of FICD under normal ER conditions has not been identified. Together, these intriguing observations raise the question of whether displacement of the autoinhibitory glutamate is the sole mode of regulation of glutamate-containing AMPylating FIC proteins, and call for investigation of diffusible signals directly able to release autoinhibition in glutamate-bearing AMPylating FIC proteins and to regulate deAMPylation in FICD.

In this study, we addressed this question by combining structural analysis and PTM assays of a single-domain FIC protein from *Enterococcus faecalis* (EfFIC), which carries an autoinhibitory glutamate in C-terminus, and of human FICD. Enteroccoci are commensals of the gastrointestinal tract that become pathogenic outside of the gut and cause difficult-to-treat infections in the hospital due to acquisition and transmission of antibiotic resistance ^22, 23^. We discover that EfFIC has both AMPylation and deAMPylation activities borne by the same active site. Furthermore, the conserved glutamate implements a Ca^2+^-controlled metal switch that tunes the balance between these activities without conformational change. Finally, we show that Ca^2+^ is an inhibitor of deAMPylation of BiP by human FICD. Our findings predict that the presence of a glutamate is a signature for AMPylation/deAMPylation bifunctionality in FIC proteins from bacteria to human, and they identify metals as diffusible signals that can directly modulate the activity of glutamate-bearing FIC proteins by binding into the active site. Importantly, they suggest that FICD is an enzymatic Ca^2+^ sensor which responds to the drop in Ca^2+^ concentration, a hallmark of ER stress, with implications in diseases that ilvolve the UPR and for therapeutic strategies.

## Results

### EfFIC is an AMPylator

*Enterococcus faecalis* FIC belongs to class III FIC proteins, which are comprised of a single FIC domain and carry an autoinhibitory glutamate in their C-terminal α-helix. We determined crystal structures of unbound, phosphate-bound, AMP-bound and ATPγS-bound wild-type EfFIC (EfFIC^WT^) and of unbound and sulfate-bound EfFIC carrying a mutation of the catalytic histidine into an alanine (EfFIC^H111A^) (**Table 1 and Table S1**). These structures were obtained in different space groups, yielding 32 independent copies of the EfFIC monomer with various environments in the crystal. All EfFIC monomers resemble closely to each other and to structures of other class III FIC proteins (**Figure 1A**). Notably, the C-terminal α-helix that bears the inhibitory glutamate shows no tendency for structural flexibility, even in subunits that are free of intersubunit contacts in the crystal. The glutamate has the same conformation as in other glutamate-bearing FIC protein structures (**Figure 1B**) and is stabilized by intramolecular interactions and, when present, by interactions with the nucleotide cofactor (**Figure 1C**). Two crystal structures were obtained in co-crystallization with a non-hydrolyzable ATP analog (ATPγS), for which well-defined electron density was observed for the ADP moiety (**Figure S1A**). The positions of the α and β phosphates of ATPγS in these structures depart markedly from those seen in ATP bound unproductively to wild type NmFIC ^7^, or bound non-canonically to CdFIC ^11^ (**Figure 1D**). In contrast, they superpose well to cofactors bound in a position competent for PTM transfer ^7,8^ (**Figure 1E**). This observation prompted us to assess whether EfFIC is competent for AMPylation, using autoAMPylation as a convenient proxy in the absence of a known physiological target (reviewed in ^20^). Using [α-^32^P]-ATP and autoradiography to measure the formation of AMPylated EfFIC (denoted ^AMP*^EfFIC^WT^), we observed that EfFIC^WT^ has conspicuous autoAMPylation activity in the presence of Mg^2+^ (**Figure 1F**). Mutation of the inhibitory glutamate into glycine (E190G) increased AMPylation, indicating that AMPylation of EfFIC^WT^ is not optimal (**Figure 1F**). We conclude from these experiments that wild-type EfFIC has canonical features of an AMPylating FIC enzyme, and that the inhibitory glutamate mitigates this activity.

**Table 1.**
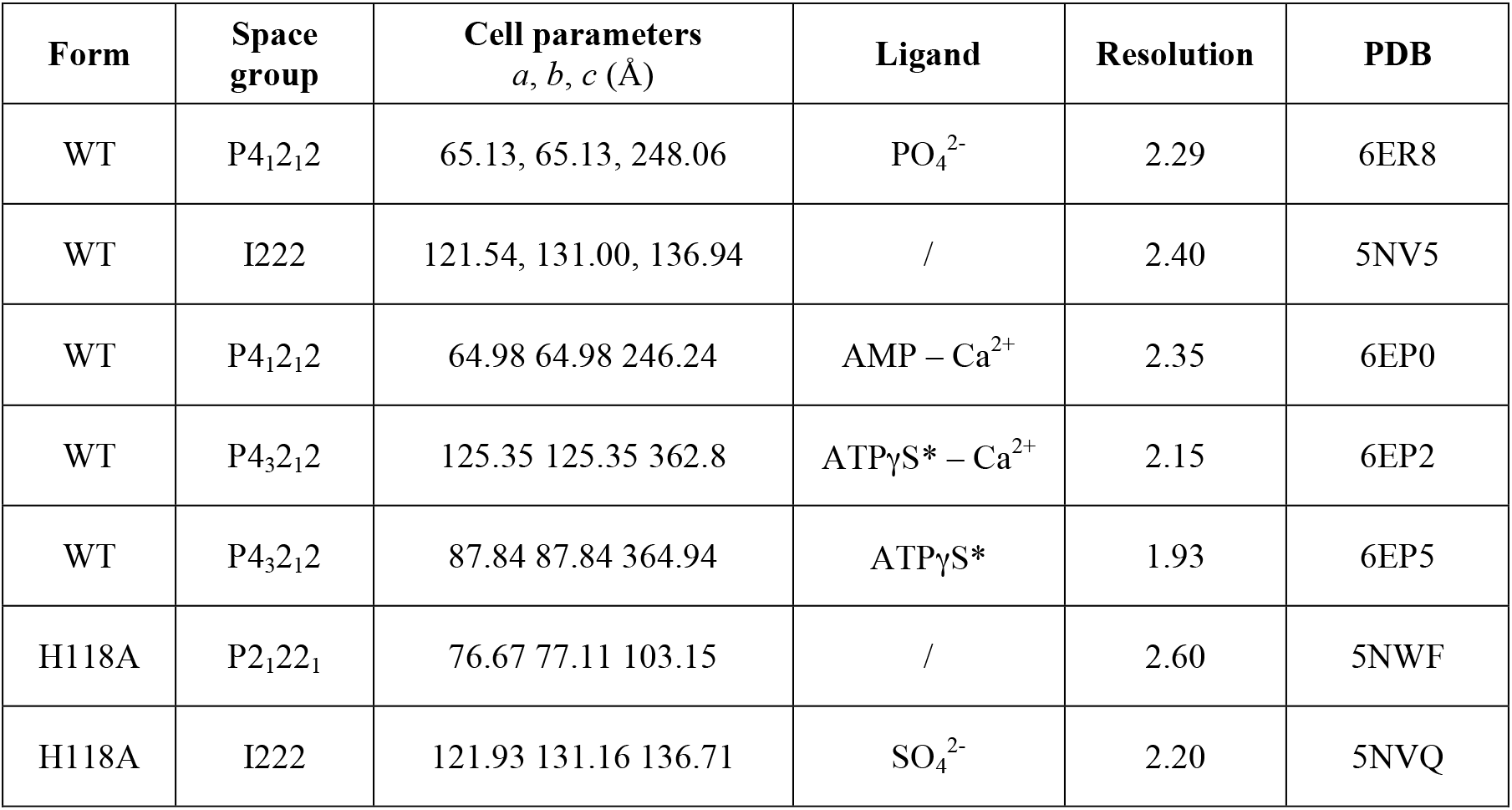
Summary of crystal structures of EfFIC determined in this study. All crystals have cell angles of α β and γ = 90°. Crystallographic statistics are given in **Table S1**. *: only the ADP moiety is visible.

| Form | Space group | Cell parameters<br>*a, b, c* (Å) | Ligand | Resolution | PDB |
| --- | --- | --- | --- | --- | --- |
| WT | P4<sub>1</sub>2<sub>1</sub>2 | 65.13, 65.13, 248.06 | PO<sub>4</sub><sup>2-</sup> | 2.29 | 6ER8 |
| WT | I222 | 121.54, 131.00, 136.94 | / | 2.40 | 5NV5 |
| WT | P4<sub>1</sub>2<sub>1</sub>2 | 64.98 64.98 246.24 | AMP – Ca<sup>2+</sup> | 2.35 | 6EP0 |
| WT | P4<sub>3</sub>2<sub>1</sub>2 | 125.35 125.35 362.8 | ATPγS* – Ca<sup>2+</sup> | 2.15 | 6EP2 |
| WT | P4<sub>3</sub>2<sub>1</sub>2 | 87.84 87.84 364.94 | ATPγS* | 1.93 | 6EP5 |
| H118A | P2<sub>1</sub>22<sub>1</sub> | 76.67 77.11 103.15 | / | 2.60 | 5NWF |
| H118A | I222 | 121.93 131.16 136.71 | SO<sub>4</sub><sup>2-</sup> | 2.20 | 5NVQ |

**Figure 1:**
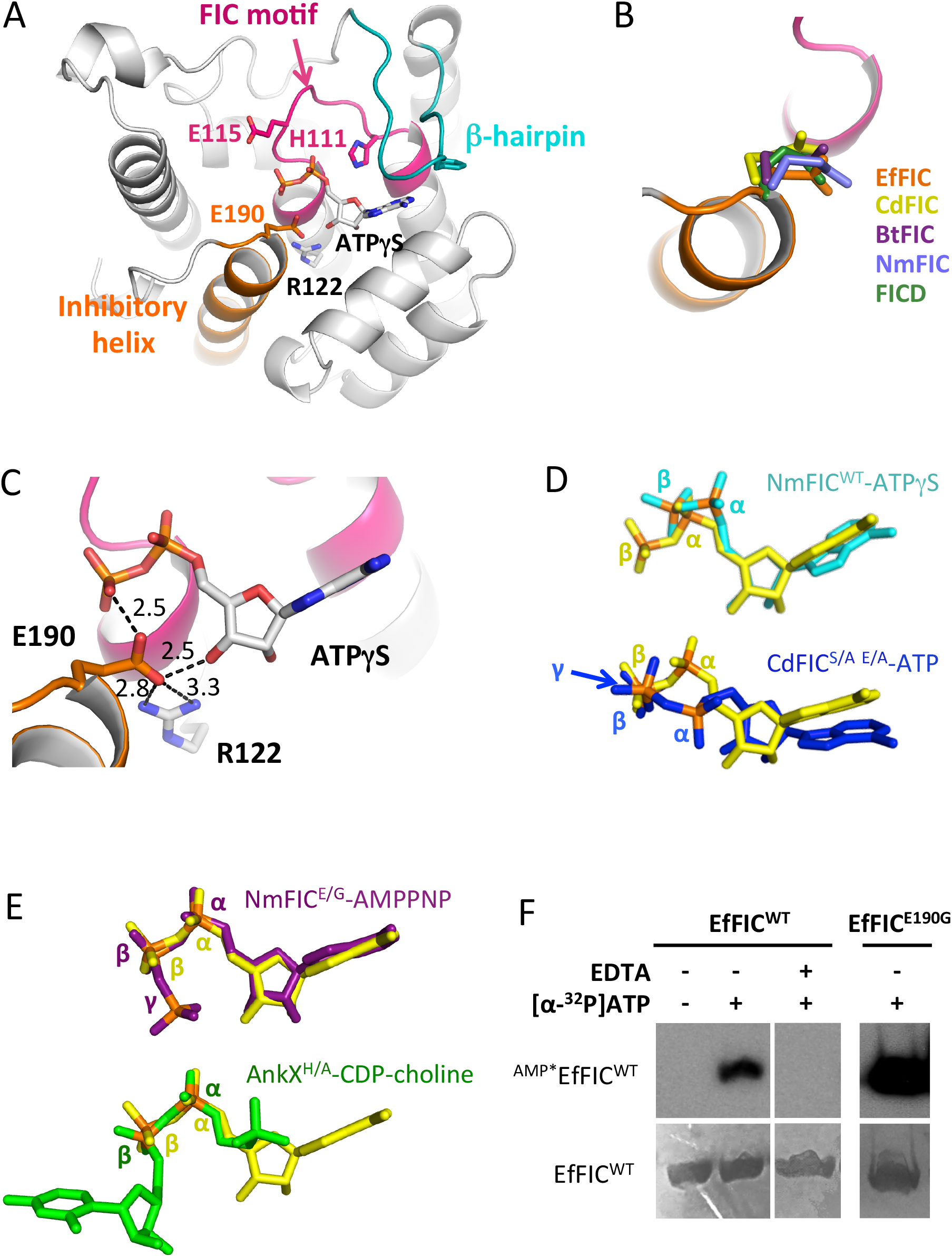
Structural basis for the AMPylation activty of EfFIC. A: Structure of the EfFIC monomer showing the FIC motif (pink), the C-terminal α-helix bearing the inhibitory glutamate (orange) and the β-hairpin predicted to bind protein substrates (cyan). The ADP moiety of ATPγS is shown in sticks. B: The inhibitory glutamate from EfFIC^WT^ (orange) is structurally equivalent to the glutamate found in the C-terminus of NmFIC (^7^, PDB 2G03) and in the N-terminus of *Bacteroides* BtFIC (PDB 3CUC), *Clostridium* CdFIC (^11^, PDB 4X2E) and of human FICD/HYPE (^10^, PDB 4U04). Superpositions are done on the structurally highly conserved FIC motif. C: Interactions of the inhibitory glutamate with the active site of ATPγS-bound EfFIC^WT^. Hydrogen bonds are depicted by dotted lines. D: The positions of the α- and β-phosphates of ATPγS bound to EfFIC^WT^ (yellow) diverge from those of ATPγS bound to NmFIC^WT^ (cyan, ^7^, PDB 3S6A) and of ATP bound to CdFIC (blue, ^11^, S31A/E35A mutant, PDB 4X2D) in a non-canonical conformations. Note that only the ADP moiety of ATPγS is visible in the EfFIC^WT^ and NmFIC^WT^ crystal structures. E: The α- and β-phosphates of ATPγS bound to EfFIC^WT^ (yellow) superpose well to those of ATP bound to NmFIC (purple, ^7^, E186G mutant, PDB 3ZLM) and of CDP-choline bound to AnkX (green, ^8^, H229A mutant, PDB 4BET) in a PTM-competent conformation. F: AutoAMPylation of EfFIC^WT^ and EfFIC^E190G^. The level of AMPylated proteins (indicated as ^AMP*^EfFIC^WT^) was measured by autoradiography using radioactive [α-^32^P]-ATP in the presence of 100 μM Mg^2+^. The reaction was carried out for one hour for EfFIC^WT^ and five minutes for EfFIC^E190G^. The total amount of EfFIC^WT^ measured by Coomassie staining in the same sample is shown.

### EfFIC is a deAMPylator in the presence of Ca^2+^

To gain further insight into the enzymatic properties of EfFIC, we solved the crystal structure of EfFIC bound to AMP (EfFIC^WT^-AMP) (**Table 1 and Table S1**). AMP superposes exactly to the AMP moiety of AMPylated CDC42 in complex with the FIC2 domain of the IbpA toxin ^5^ (**Figure 2A)**. Electron-rich density was observed next to AMP in the active site, corresponding to a calcium ion present in the crystallization solution to the exclusion of all other metal ions (**Figure S1B**). Ca^2+^ has 6 coordinations with distances in the expected 2.1-2.9 Å range, arranged with heptahedral geometry in which one ligand, which would be located opposite to one phosphate oxygen, is missing. It interacts with the phosphate of AMP, with the acidic residue in the FIC motif (Glu115), and with the inhibitory glutamate (Glu190) through a water molecule (**Figure 2B**). The position of Ca^2+^ in the EfFIC^WT^-AMP structure differs from that of Mg^2+^ observed in other FIC protein structures in complex with ATP (**Figure 2C**), raising the intriguing issue that it may have an alternative catalytic function. Inspired by the recent observation that animal FICD proteins have deAMPylation enzymatic activity ^18, 24^, we analyzed whether EfFIC would have deAMPylation activity in the presence of Ca^2+^. Remarkably, addition of Ca^2+^ to EfFIC^WT^ that had been previously autoAMPylated in the presence of Mg^2+^ and [α-^32^P]-ATP induced conspicuous deAMPylation (**Figure 2D**).

**Figure 2:**
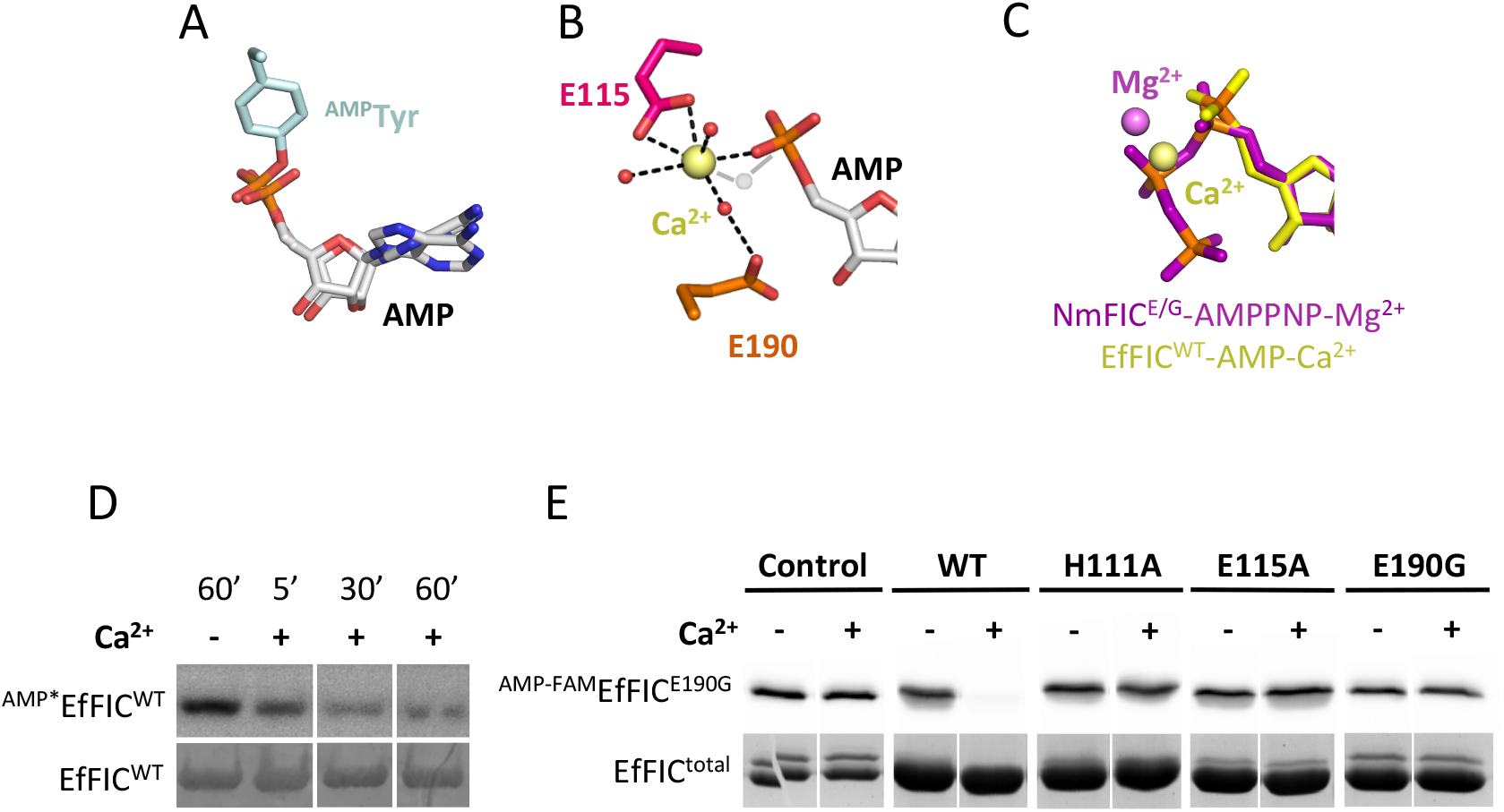
EfFIC is a deAMPylator in the presence of Ca^2+^. A: AMP bound to EfFIC^WT^ superposes with the AMP moiety of AMPylated CDC42 in complex with the FIC protein IbpA (^5^, PDB 4ITR). Superposition was carried out based on the FIC motif. B: Close-up view of Ca^2+^ bound to EfFIC^WT^-AMP. The incomplete heptagonal coordination sphere of Ca^2+^ is indicated by dotted lines. Water molecules are shown as red spheres. The proposed position of the water molecule responsible for in-line nucleophilic attack is indicated in grey. C: Ca^2+^ bound to EfFIC^WT^-AMP is shifted by 1.3 Å with respect to Mg^2+^ bound to NmFIC^E/G^-AMPPNP. The superposition is done on the FIC motif. D: EfFIC^WT^ has deAMPylation activity in the presence of Ca^2+^. EfFIC^WT^ was autoAMPylated with [α-^32^P]-ATP in the presence 100 nM Mg^2+^ for one hour, then the sample was incubated for 60 minutes with EDTA alone (1mM) (lane “-“) or 5, 30 and 60 minutes with EDTA (1mM) and an excess of Ca^2+^ (10 mM) (lanes “+”). AMPylation levels were analyzed by autoradiography (upper panel). The total amount of EfFIC^WT^ in each sample was measured by Coomassie staining (lower panel). E: Key residues of the AMPylation active site are required for deAMPylation. EfFIC^E190G^ was auto-AMPylated for one hour in the presence of fluorescently-labeled ATP-FAM and Mg^2+^, then ^AMP-FAM^EfFIC^E190G^ was purified to remove Mg^2+^, PPi and ATP-FAM. DeAMPylation was then triggered by addition of wild-type or mutant EfFIC as indicated in the presence or not of 1mM Ca^2+^. AMPylation levels (indicated as ^AMP-FAM^EfFIC^E190G^) after one hour incubation were analyzed by fluorescence (upper panel). The total amount of EfFIC proteins (indicated as EfFIC^total^) in each sample was measured by Coomassie staining (lower panel).

In the above setup, the AMPylation and deAMPylation activities are potentially acting concurrently. To characterize the deAMPylation reaction selectively, the hyperactive EfFIC^E190G^ mutant was autoAMPylated in the presence of Mg^2+^, purified to remove ATP, PPi and Mg^2+^ such that no AMPylation remains possible, then its deAMPylation was triggered by addition of EfFIC^WT^ or EfFIC mutants in the presence of Ca^2+^. The level of AMPylated EfFIC (denoted ^AMP-FAM^EfFIC) was quantified by fluorescence using ATP-FAM, an ATP analog fluorescently labeled on the adenine base. Robust deAMPylation was observed upon addition of EfFIC^WT^ and Ca^2+^ (**Figure 2E**, EfFIC^WT^ panel). No spontaneous deAMPylation of ^AMP-FAM^EfFIC^E190G^ was observed in the absence of EfFIC^WT^ (**Figure 2E**, control panel), indicating that the deAMPylation reaction occurs in trans. We used this deAMPylation setup to identify residues critical for deAMPylation (**Figure 2E**, mutant panels). Mutation of the catalytic histidine (H111A) and of the metal-binding acidic residue in the FIC motif (E115A) impaired deAMPylation of ^AMP-FAM^EfFIC^E190G^. Likewise, EfFIC^E190G^, which carries the mutation of the inhibitory glutamate, was unable to catalyze deAMPylation, consistent with the absence of spontaneous deAMPylation in the assay. We conclude from these experiments that EfFIC is a bifunctional enzyme, that AMPylation and deAMPylation are borne by the same active site, and that the inhibitory glutamate is involved in the deAMPylation reaction.

### The AMPylation and deAMPylation reactions are differentially regulated by metals

The above results raise the issue of the nature of signals acting on the bifunctional active site of EfFIC to regulate AMPylation/deAMPylation alternation. Previous work showed that AMPylation of *Escherichia coli* DNA gyrase by NmFIC, which shares 56% sequence identity with EfFIC, was highly sensitive to the toxin concentration, with a sharp drop of activity above 250 µM ^21^. We used purified ^AMP-FAM^EfFIC^E190G^ to analyze whether the deAMPylation activity of EfFIC^WT^ would be similarly inhibited by increasing concentrations of EfFIC^WT^ (1-2000 nM). As shown in **Figure 3A**, deAMPylation increased with EfFIC^WT^ concentration, indicating that this reaction is not adversely affected by EfFIC concentration. Alternatively, we reasoned that the distinct electrochemical properties of Ca^2+^ and Mg^2+^ (reviewed in ^25^) may allow them to support AMPylation and deAMPylation differentially. Remarkably, Ca^2+^ was unable to support AMPylation, contrary to Mg^2+^ (**Figure 3B**). In contrast, both Mg^2+^ and Ca^2+^ supported potent deAMPylation (**Figure 3C, left panel**). Importantly, mutation of the inhibitory glutamate eliminated the ability of EfFIC to use Ca^2+^ for deAMPylation, while the mutant retained partial deAMPylation in the presence of Mg^2+^ (**Figure 3C, right panel**). To understand how Ca^2+^ affects AMPylation and deAMPylation differentially, we determined the crystal structure of EfFIC^WT^-ATPγS-Ca^2+^. In this structure, Ca^2+^ is heptacoordinated to the α- and β-phosphates of ATPγS (of which again only the ADP moiety is visible), to the inhibitory glutamate and to 4 water molecules (**Figure 3D**). Strikingly, Ca^2+^ is shifted by 3.2 Å from Mg^2+^ bound to NmFIC in an equivalent configuration ^7^ (**Figure 3E**) and it lacks an interaction with the acidic residue from the FIC motif, which explains why it inhibits AMPylation. Together, these observations point towards a 3-position metal switch differentially operated by Mg^2+^ and Ca^2+^, which includes a position competent for AMPylation, a position that inhibits AMPylation and a position competent for deAMPylation. To test this hypothesis, we measured the apparent AMPylation efficiency of EfFIC^WT^ at different Mg^2+^/Ca^2+^ ratio. As shown in **Figure 3F**, AMPylation is prominent when Mg^2+^ exceeds Ca^2+^, while Ca^2+^ in excess over Mg^2+^ favors deAMPylation. We conclude from these experiments that competition between Mg^2+^ and Ca^2+^ regulates the balance between AMPylation and deAMPylation efficiencies and that this regulatory metal switch is implemented by differential usage of the inhibitory glutamate and the acidic residue in the FIC motif for binding metals.

**Figure 3:**
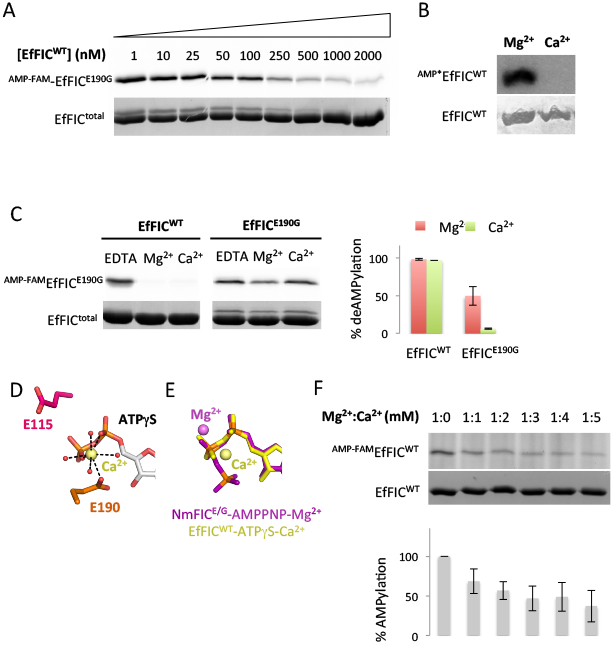
The Mg^2+^/Ca^2+^ ratio tunes the balance between EfFIC AMPylation and de-AMPylation activities. A: DeAMPylation is not inhibited at high EfFIC concentration. Purified ^AMP-FAM^EfFIC^E190G^ was incubated with increasing concentrations of EfFIC^WT^ for one hour in the presence of 100µM Ca^2+^. AMPylation levels were measured by fluorescence as in **Figure 2D**. B: EfFIC^WT^ uses Mg^2+^ but not Ca^2+^ for autoAMPylation. AutoAMPylation was carried out at 1mM Ca^2+^ or Mg^2+^ for one hour and was measured by autoradiography as in **Figure 1F**. C: The conserved glutamate of EfFIC is required for the usage of both Mg^2+^ and Ca^2+^ for deAMPylation. Experiments were carried out for one hour as in **Figure 3A** in the presence of 0.5mM EDTA and 3mM of Mg^2+^ or Ca^2+^. Quantification of deAMPylation efficiencies is shown on the right panel. D: Close-up view of Ca^2+^ bound to EfFIC^WT^-ATPγS. The heptagonal coordination sphere of Ca^2+^ is indicated by dotted lines. Water molecules are shown as red spheres. E: Ca^2+^ bound to EfFIC^WT^-ATPγS is shifted with respect to Mg^2+^ bound to AMPylation-competent NmFIC^E/G^-AMPPNP (PDF 2ZLM). F: The net AMPylation efficiency of EfFIC^WT^ is controled by the Mg^2+^/Ca^2+^ ratio. EfFIC^WT^ was incubated for one hour in the presence of ATP-FAM at fixed Mg^2+^ concentration (1mM) and increasing concentrations of Ca^2+^. ^AMP-FAM^EfFIC levels were measured by fluorescence. The steady-state AMPylation efficiency, which results from the relative AMPylation and deAMPylation rates, is expressed as the percentage of the maximal AMPylation level obtained with Mg^2+^ alone. All measurements have p-values <0.05 with respect to the experiment containing Mg^2+^ alone. The total amount of EfFIC^WT^ measured by Coomassie staining in each sample is shown.

### Ca^2+^ tunes deAMPylation of the BiP chaperone by human FICD

DeAMPylation of the BiP chaperone has been recently identified as the primary in vitro activity of human FICD ^18^, which features a glutamate structurally equivalent to the inhibitory glutamate in EfFIC (see **Figure 1B**, ^10^) that is critical for deAMPylation ^18^. We analyzed whether, as observed in EfFIC, Mg^2+^ and Ca^2+^ metals could also affect FICD activity, using fluorescent ATP-FAM to monitor BiP AMPylation. No measurable AMPylation of BiP by FICD^WT^ was observed, neither with Mg^2+^ nor Ca^2+^, although FICD^WT^ itself showed weak autoAMPylation in the presence of both metals (**Figure 4A**). Alternatively, we used FICD^E234G^, in which the conserved glutamate is mutated to glycine, to produce AMPylated BiP. Remarkably, while purified ^AMP-FAM^BIP was efficiently deAMPylated by FICD^WT^ in the presence of Mg^2+^, no deAMPylation was measured in the presence of Ca^2+^ (**Figure 4B**). To determine whether FICD does not bind Ca^2+^ or is unable to use it for deAMPylation, we carried out an Mg^2+^/Ca^2+^ competition experiment in which FICD^WT^ and purified ^AMP-FAM^BiP were incubated at increasing Ca^2+^ concentration and a fixed Mg^2+^ concentration. As shown in **Figure 4C**, deAMPylation efficiency decreased as the Ca^2+^/Mg^2+^ ratio increased, suggesting that Ca^2+^ inhibits deAMPylation by competing with Mg^2+^. We conclude from these experiments that Ca^2+^ binds to FICD in an inhibitory manner, which allows it to tune the deAMPylation efficiency of FICD towards the BiP chaperone.

**Figure 4:**
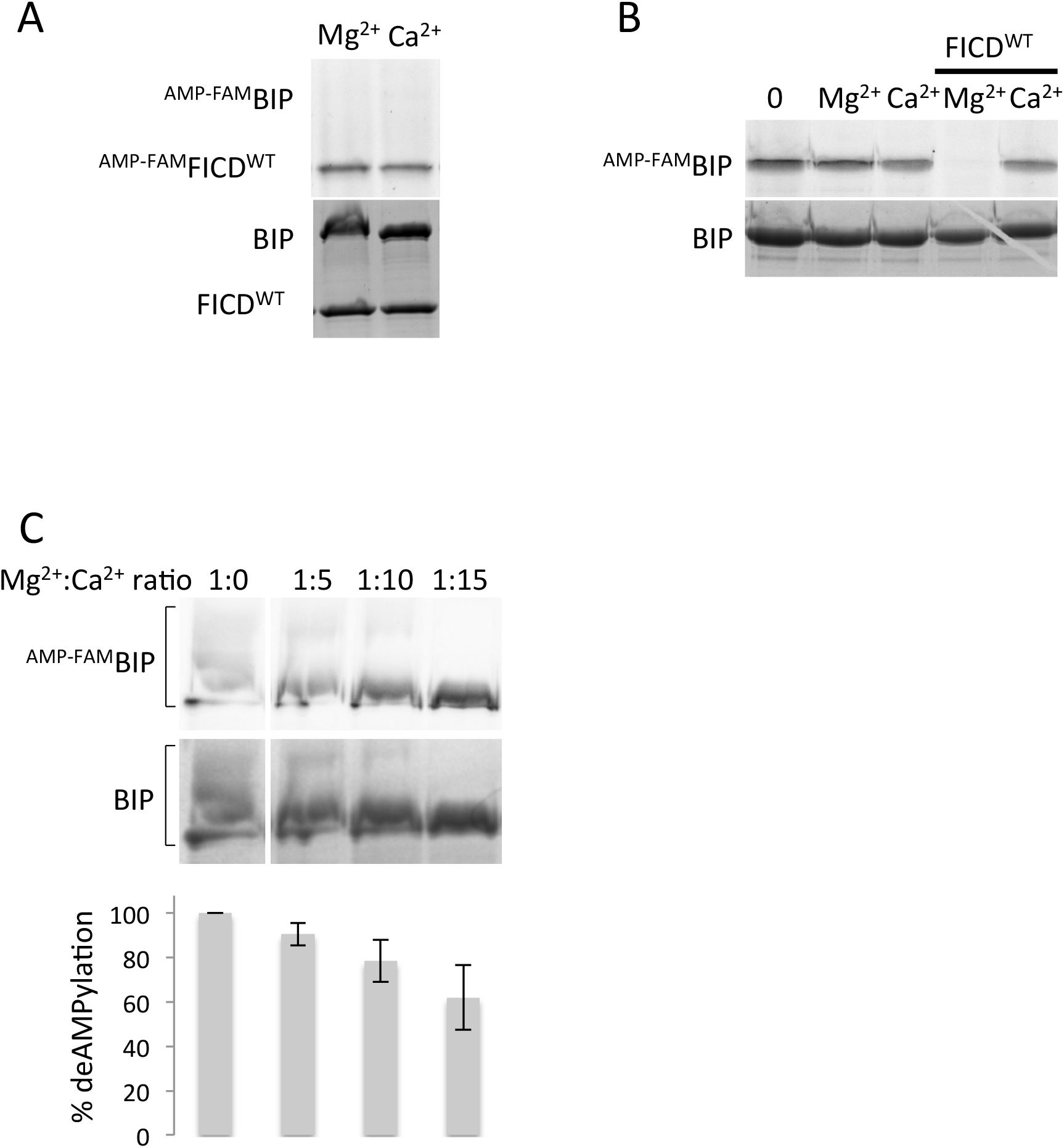
DeAMPylation of BiP by FICD requires Mg^2+^ and is inhibited by Ca^2+^. A: FICD^WT^ does not AMPylate the BiP chaperone. Reactions were carried out for one hour in presence of 1mM Mg^2+^ or 1mM Ca^2+^. The level of AMPylated proteins (indicated as ^AMP-FAM^FICD^WT^ and ^AMP-FAM^BIP) was measured by fluorescence. The total amount of proteins measured by Coomassie staining in the same sample is shown. B: Ca^2+^ inhibits deAMPylation of BiP by FICD/HYPE^WT^. BiP was first AMPylated for one hour by the hyperactive FICD^E234G^ mutant in the presence of 100 μM Mg^2+^, then ^AMP-FAM^BIP was purified to remove Mg^2+^, PPi and ATP-FAM. DeAMPylation of ^AMP-FAM^BIP was then triggered by addition of wild-type or mutant FICD as indicated in the presence of 1mM Mg^2+^ or 1mM Ca^2+^. ^AMP-FAM^BiP levels were measured by fluorescence. The total amount of BiP measured by Coomassie staining in each sample is shown. C: The net deAMPylation efficiency of FICD^WT^ is tuned by the Mg^2+^/Ca^2+^ ratio. DeAMPylation was carried out as in Figure 4B using a fixed Mg^2+^ concentration (200µM) and 0, 1, 2 or 3mM Ca^2+^. AMPylation levels were measured by fluorescence, normalized to the fluorescence intensity of FICD and expressed as the percentage of maximal deAMPylation level obtained in the absence of Ca^2+^. All data have p-values <0.05 with respect to the control in the absence of Ca^2+^. The total amount of BiP measured by Coomassie staining in each sample is shown.

## Discussion

In this study, we sought after a diffusible signal able to regulate directly the large group of AMPylating FIC proteins in which the AMPylation activity is repressed by a conserved glutamate within the active site. Combining crystallography and PTM assays, we first show that bacterial EfFIC is a bifunctional enzyme that encodes AMPylating and deAMPylating activities and that both reactions use the same active site. Next, we discover that the balance between these opposing activities is controlled by a metal switch, in which each reaction is differentially supported and inhibited by Mg^2+^ and Ca^2+^ in a manner that the Mg^2+^/Ca^2+^ ratio determines the net AMPylation level. Furthermore, we identify the inhibitory glutamate and the acidic residue in the FIC motif as residues essential for the metal switch. Finally, we show that deAMPylation of the endoplasmic reticulum BiP chaperone by human FICD is also dependent on the Ca^2+^/Mg^2+^ ratio, with high Ca^2+^ concentration inhibiting deAMPylation.

The identification of a potent deAMPylation activity in a bacterial FIC protein (this study) and in human FICD ^18^, which depends on a structurally equivalent glutamate in these otherwise remotely related FIC proteins, leads us to propose that the conserved glutamate is a signature of the ability of FIC proteins to catalyze both AMPylation and deAMPylation. Our data allow to delineate the catalytic basis for this bifunctionality, in which catalytic residues are shared by the AMPylation and deAMPylation reactions but have different roles in catalysis. In the AMPylation reaction, the invariant histidine in the FIC motif activates the acceptor hydroxyl of a target protein by attracting a proton and the acidic residue (Asp or Glu) in the FIC motif binds a metal that stabilizes the phosphates of the ATP cofactor at the transition state (reviewed in ^2, 3^). Based on the observations that the α- and β-phosphates of ATP bind with canonical positions in wild-type EfFIC and that AMPylation is potentiated by mutation of the glutamate in various FIC proteins, we propose that the primary role of the glutamate in AMPylation is to mitigate the efficiency of this reaction in the presence of Mg^2+^, rather than to fully repress it, possibly in order to match AMPylation and deAMPylation efficiencies. In the deAMPylation mechanism depicted in **Figure 5** (see also discussion in **Supplementary data and Figure S2**), the conserved glutamate activates a water molecule for nucleophilic attack of the phosphorus of the AMP moiety, and the invariant histidine generates the free hydroxyl group in the protein residue by giving up a proton, as also proposed in ^18^. The nucleophilic water molecule is readily identified as the missing ligand in the coordination sphere of Ca^2+^ in our EfFIC-AMP-Ca^2+^ structure, where it would be precisely positioned for in-line nucleophilic attack (**see Figure 2B**). Interestingly, the metal is bound to both the acidic residue of the FIC motif and the conserved glutamate in this structure, which positions it to stabilize negative charges in the AMPylated substrate that develop at the transition state.

**Figure 5:**
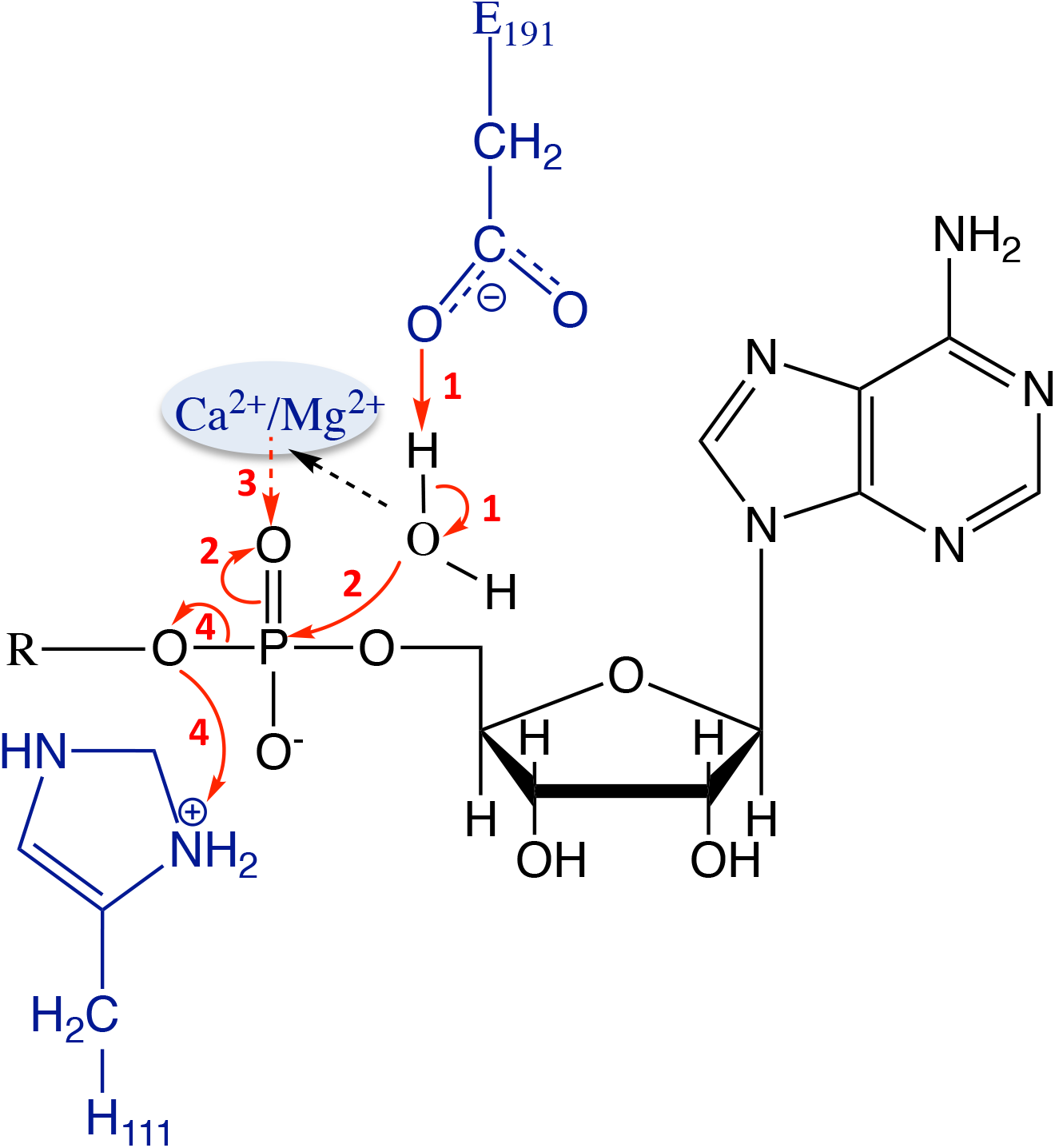
Model of the Ca^2+^-assisted deAMPylation catalytic mechanism. In this model, the regulatory glutamate (Glu190 in EfFIC) attracts a proton from a water molecule coordinating the metallic cation (1) to activate it for nucleophilic attack of its oxygen on the phosphorus of AMP moiety in the AMPylated substrate (2). The positive charge provided by Ca^2+^ increases electrophilicity of the phosphorus and stabilizes the negative charge of the intermediate (2 and 3). The intermediate harboring a pentavalent phosphorus then rearranges, leading to the breaking of the phosphor-ester bond, which is elicited by the capture of a proton provided by the catalytic histidine

Importantly, our data identify for the first time a diffusible signal able to modulate the activity of glutamate-bearing FIC proteins. A remarkable feature in the above bifunctional mechanism is that both reactions can be adversely regulated by a second metal that competes with the catalytic metal. In EfFIC, we observed that Ca^2+^ binds to ATP in a position that is shifted with respect to the canonical AMPylation Mg^2+^-binding site, resulting in decreased AMPylation. In a related scenario, Ca^2+^ competes with Mg^2+^ in the deAMPylation reaction catalyzed by FICD, thereby decreasing deAMPylation. This multi-position metal switch constitutes a new paradigm in bifunctional enzyme regulation, in which the relative affinities of specific metals for the AMPylation and deAMPylation configurations tip the balance towards opposing activities within the same active site. Future studies are now needed to determine the bifunctionality spectrum of glutamate-bearing FIC proteins resulting from variations in metal specificities and affinities. In addition, observations by us and others that FICD has distinct AMPylation and deAMPylation patterns towards itself and BiP suggests that the protein substrate influences the AMPylation/deAMPylation balance through still unknown mechanisms, which will have to be investigated. Another important question for future studies is how other levels of regulation, such as autoAMPylations and changes in oligomerization that have been described for a close homolog of EfFIC ^21^, combine with the intrinsic metal switch identified in this study.

Our findings predict that bacterial glutamate-containing FIC proteins are bifunctional enzymes able to switch between AMPylation and deAMPylation activities upon changes in metal homeostasis. An appealing hypothesis is thus that glutamate-bearing FIC proteins function as toxin/antitoxin (TA) modules, in which the toxin and antitoxin activities would be encoded within the same active site under the control of regulatory metals. They would therefore differ from type III TA modules, in which the toxin is repressed by direct interaction with the antitoxin, or type IV TA modules, in which the effects of the toxin are reverted by the antitoxin without direct interaction (reviewed in ^26^). It is interesting to note that the antitoxin component of TA modules such as Bartonella VbhT/VbhA features a structurally equivalent glutamate that points into the catalytic site of the toxin ^7^, raising the question of whether there may exist conditions where the antitoxin glutamate implements deAMPylation within the TA complex. The physiological conditions that involve variations in metals homeostasis to activate the switch between opposing activities in these FIC proteins, and how individual bacteria exploit the bifunctionality of their FIC proteins within their ecological niche or in infections will be important issues to address in future studies.

Finally, our discovery that deAMPylation of the BiP chaperone by human FICD is modulated by changes in Ca^2+^ concentration raises important questions with respect to the regulation and role of FICD in endoplasmic reticulum (ER) functions. The ER is the major organelle involved in Ca^2+^ homeostasis (reviewed in ^27-29^), and the total Ca^2+^ concentration in the ER is as high as 2-3 mM under resting conditions ^30^. Disruption of Ca^2+^ homeostasis, such as depletion of the ER Ca^2+^ store induced by the drug thapsigargin, swiftly alters protein folding processes and activates the unfolded protein response ^31^. Large Ca^2+^ fluctuations are therefore considered as major determinants of ER stress responses (reviewed in ^28,29^). On the other hand, there is currently no evidence that the Mg^2+^ concentration changes significantly in the ER over time (reviewed in ^32^). Thus, while keeping in mind that Ca^2+^ and Mg^2+^ concentrations and local gradients in the ER have remained difficult to determine with accuracy, an essentially constant Mg^2+^ concentration and Ca^2+^ fluctuations in the 10µM-3mM range, such as those used in our vitro assays, are plausibly encountered in the ER as it experiences transition from resting to stress conditions. Recently, FICD has been demonstrated to stimulate the activity of the BiP chaperone in response to an increase in the unfolded protein load ^13–15,24^, and this activation was correlated to deAMPylation of BiP by FICD ^18^. It is thus tempting to speculate that inhibition of FICD deAMPylation activity by high Ca^2+^, which we observe *in vitro*, reflects its inhibition under ER homeostasis, where Ca^2+^ concentration is high and BiP activity is low. Conversely, depletion of Ca^2+^ induces ER stress and triggers the UPR. Depletion of Ca^2+^ may thus release inhibition of FICD deAMPylation activity, leading to efficient deAMPylation of BiP and up-regulation of its activity, which is a key feature of the UPR. We propose that FICD is has features of an enzymatic sensor of Ca^2+^ in the ER, which could allow it to function as an integrator between Ca^2+^ homeostasis in the ER and the BiP-mediated unfolded protein response.

In conclusion, we have identified Ca^2+^ as a diffusable signal that modulates the intrinsic enzymatic activity of glutamate-bearing FIC proteins directly and tips the balance between AMPylation and deAMPylation reactions without a conformational change in the enzyme. Future studies are now needed to investigate how the metal switch of glutamate-bearing FIC protein activities is exploited in bacterial stress, and, in the case of animal FICD, its role in the unfolded protein response and its crosstalks with Ca^2+^-controled processes in the ER. The detrimental effects of prolonged UPR in a myriad of diseases (reviewed in ^33,34^) makes new druggable targets highly desirable (reviewed in ^35^). Given the established pivotal role of the BiP chaperone (reviewed in ^16^) and of calcium (reviewed in ^29^) in this process, the Ca^2+^-regulated activity of FICD uncovered in this study may constitute a new druggable target.

## Material and Methods

### Protein cloning, expression and purification

The codon-optimized gene encoding full-length *Enterococcus faecalis* EfFIC with an N-terminal 6-histidine tag was from GeneArt Gene Synthesis (ThermoFisher Scientific) and cloned into a pET22b(+) vector. The codon-optimized gene encoding human FICD/HYPE (residues 45-459) carrying an N-terminal 6-His tag followed by SUMO tag was from GeneArt Gene Synthesis and cloned into a pET151/D-TOPO vector (ThermoFisher Scientific). All mutations were performed with the QuickChange II mutagenesis kit (Agilent). *Mus musculus* BIP in pUJ4 plasmid is a kind gift from Ronald Melki (CNRS, Gif-sur-Yvette). All constructs were verified by sequencing (GATC). All EfFIC constructs were expressed in *E. coli* BL21 (DE3) pLysS in LB medium. Overexpression was induced overnight with 0.5 mM IPTG at 20°C. Bacterial cultures were centrifuged for 40 min at 4000g. Bacterial pellets were resuspended in lysis buffer (50 mM Tris-HCl pH 8.0, 150 mM NaCl, 5% glycerol, 0.25 mg/mL lysozyme) containing a protease inhibitor cocktail, disrupted at 125 psi using a high pressure cell disrupter and centrifuged 30 min at 22000g. The cleared lysate supernatant was loaded on a Ni-NTA affinity chromatography column (HisTrap FF, GE Healthcare) and eluted with 250 mM imidazole. Purification was polished by gel filtration on a Superdex 200 16/600 column (GE Healthcare) equilibrated with storage buffer (50 mM Tris-HCl pH 8.0, 150 mM NaCl). Wild-type and mutant FICD/HYPE were expressed and purified as EfFIC, except that the lysis buffer was complemented with 1mM DTT and 0.02% Triton X-100 and other buffers with 1mM DTT. To remove the SUMO tag, FICD/HYPE was incubated with SUMO protease (ThermoFischer) at 1/100 weight/weight ratio during 1 hour at room temperature. The cleaved fraction was separated by affinity chromatography (HisTrap FF, GE Healthcare) and further purified by gel filtration on a Superdex 200 10/300 column (GE Healthcare) equilibrated with storage buffer (50 mM Tris pH 8.0, 150 mM NaCl, 1mM DTT, 5% glycerol). Mouse BIP was expressed and purified essentially as EfFIC.

### Crystallization and structure determination

A summary of the crystal structures determined in this study is in **Table 1**. Proteins were crystallized using a TTP Labtech’s Mosquito LCP crystallization robot and crystallization screens (Jena Bioscience and Quiagen). Conditions leading to crystals were subsequently optimized. Diffraction data sets were recorded at synchrotron SOLEIL and ESRF. Datasets were processed using XDS ^36^, xdsme (https://github.com/legrandp/xdsme) or autoProc ^37^. Structures were solved by molecular replacement and refined with the Phenix suite ^38^ or Buster (Bricogne G., Blanc E., Brandl M., Flensburg C., Keller P., Paciorek W., Roversi P, Sharff A., Smart O.S., Vonrhein C., Womack T.O. (2017). BUSTER version 2.10.2. Cambridge, United Kingdom: Global Phasing Ltd.). Models were build using Coot ^39^. Softwares used in this project were curated by SBGrid ^40^. Crystallization conditions, data collection statistics and refinement statistics are given in **Table S1**. All structures have been deposited with the Protein Data Bank (PDB codes in **Table S1**).

### AMPylation and deAMPylation assays

AMPylation and deAMPylation autoradiography assays were carried out using the following protocols. For AMPylation reactions, 8 µg of purified proteins were mixed with 10 µCi [α-^32^P] ATP (Perkin Elmer) in a buffer containing 50mM Tris-HCl pH 7.4, 150 mM NaCl and 0.1 mM MgCl_2_. Reactions were incubated for 1 h at 30 °C, then stopped with reducing SDS sample buffer and boiling for 5 min. For deAMPylation, AMPylation was performed as above, then 1 mM EDTA was added with or without 10 mM CaCl_2_. Proteins were resolved by SDS-PAGE and AMPylation was revealed by autoradiography. EfFIC AMPylation and deAMPylation fluorescence assays were carried out using the following protocols. AMPylation was carried out using a fluorescent ATP analog modified by N^6^-(6-Amino)hexyl on the adenine base (ATP-FAM, Jena Bioscience). AMPylated proteins were obtained by incubation for one hour at 30 °C in 50mM Tris pH 8.0, 150 mM NaCl, 0.1 mM MgCl_2_ and an equimolar amount of ATP-FAM. Before deAMPylation reactions, the buffer was exchanged to 50 mM Tris-HCl pH 8.0 and 150 mM NaCl by 5 cycles of dilution/concentration on a Vivaspin-500 with a cut-off of 10kDa (Sartorius), resulting in a final dilution of ATP-FAM, MgCl_2_ and PPi by about 10^5^ times. DeAMPylation reactions were carried out using 3 µM of AMPylated protein and 6 µM of freshly purified EfFIC proteins (except for the experiment depicted in Figure 3A) in a buffer containing 50mM Tris-HCl pH 8.0 and 150 mM NaCl, for 1h at 30°C. Reactions were stopped by addition of reducing SDS sample buffer and boiling for 5 min. Proteins were resolved by SDS-PAGE and modification by AMP-FAM was revealed by fluorescence using green channel (excitation: 488 nm, emission: 526 nm) on a Chemidoc XR+ Imaging System (BioRad). FICD AMPylation and deAMPylation fluorescence assays were carried out using the following protocols. AMPylation was carried out using fluorescent ATP-FAM. AMPylated BIP was obtained by incubation for one hour at 30 °C in 50mM Tris pH 8.0, 150 mM NaCl, 0.1 mM MgCl_2_, 2µM FICD^E234G^ and an equimolar amount of ATP-FAM. Before deAMPylation reactions, the buffer was exchanged with 50mM Tris-HCl pH 8.0 and 150 mM NaCl by 5 cycles of dilution/concentration on a Vivaspin-500 with a cut-off of 50kDa (Sartorius), resulting in a final dilution of ATP-FAM, MgCl_2_ and PPi produced by the reaction of about 10^5^ times. DeAMPylation reactions were carried using 2 µg of AMPylated protein and 4 µg of freshly purified FICD^WT^ in a buffer containing 50mM Tris pH 8.0 and 150 mM NaCl, for 1h at 30°C. Reactions were stopped and ATP-FAM modification revealed as EfFIC. Quantification of AMP-FAM levels was done using (ImageLab, BioRad). All experiments were done at least in triplicate, except kinetics in Figure 3A that were done in duplicate.

## Acknowledgements

This work was supported by grants from the Fondation pour la Recherche Médicale and from the Agence Nationale pour la Recherche to J.C. and by grants n°ANR-10-LABX-62-IBEID and from the Fondation pour la Recherche Médicale to C.B. S.V. was supported by a PhD grant from the DIM MALINF and G.O. by a stipend from the Pasteur-Paris University International PhD program. We are grateful to the scientific teams at the PX1 and PX2 beamlines at the SOLEIL synchrotron (Gif-sur-Yvette, France) and from the ID29, ID30-A3 and ID30B beamlines at the European Synchrotron Research Facility (ESRF, Grenoble, France) for their expertise and advice. We thank Pascale Serror (INRA, Jouy-en-Josas, France) and Philippe Glaser (Institut Pasteur) for discussions and the members of the Cherfils lab for help and shared expertise.

## References

1. Kinch, L.N., Yarbrough, M.L., Orth, K. & Grishin, N.V. Fido, a novel AMPylation domain common to fic, doc, and AvrB. PLoS One 4, e5818 (2009).

2. Garcia-Pino, A., Zenkin, N. & Loris, R. The many faces of Fic: structural and functional aspects of Fic enzymes. Trends Biochem Sci 39, 121–9 (2014).

3. Roy, C.R. & Cherfils, J. Structure and function of Fic proteins. Nat Rev Microbiol 13, 631–40 (2015).

4. Harms, A., Stanger, F.V. & Dehio, C. Biological Diversity and Molecular Plasticity of FIC Domain Proteins. Annu Rev Microbiol 70, 341–60 (2016).

5. Xiao, J., Worby, C.A., Mattoo, S., Sankaran, B. & Dixon, J.E. Structural basis of Fic-mediated adenylylation. Nat Struct Mol Biol 17, 1004–10 (2010).

6. Luong, P. et al. Kinetic and structural insights into the mechanism of AMPylation by VopS Fic domain. J Biol Chem 285, 20155–63 (2010).

7. Engel, P. et al. Adenylylation control by intra- or intermolecular active-site obstruction in Fic proteins. Nature 482, 107–10 (2012).

8. Campanacci, V., Mukherjee, S., Roy, C.R. & Cherfils, J. Structure of the Legionella effector AnkX reveals the mechanism of phosphocholine transfer by the FIC domain. EMBO J 32, 1469–77 (2013).

9. Castro-Roa, D. et al. The Fic protein Doc uses an inverted substrate to phosphorylate and inactivate EF-Tu. Nat Chem Biol 9, 811–7 (2013).

10. Bunney, T.D. et al. Crystal structure of the human, FIC-domain containing protein HYPE and implications for its functions. Structure 22, 1831–43 (2014).

11. Dedic, E. et al. A Novel Fic (Filamentation Induced by cAMP) Protein from Clostridium difficile Reveals an Inhibitory Motif-independent Adenylylation/AMPylation Mechanism. J Biol Chem 291, 13286–300 (2016).

12. Yarbrough, M.L. et al. AMPylation of Rho GTPases by Vibrio VopS disrupts effector binding and downstream signaling. Science 323, 269–72 (2009).

13. Sanyal, A. et al. A novel link between Fic (filamentation induced by cAMP)-mediated adenylylation/AMPylation and the unfolded protein response. J Biol Chem 290, 8482–99 (2015).

14. Ham, H. et al. Unfolded protein response-regulated Drosophila Fic (dFic) protein reversibly AMPylates BiP chaperone during endoplasmic reticulum homeostasis. J Biol Chem 289, 36059–69 (2014).

15. Preissler, S. et al. AMPylation matches BiP activity to client protein load in the endoplasmic reticulum. Elife 4, e12621 (2015).

16. Zhao, L. & Ackerman, S.L. Endoplasmic reticulum stress in health and disease. Curr Opin Cell Biol 18, 444–52 (2006).

17. Preissler, S. et al. AMPylation targets the rate-limiting step of BiP’s ATPase cycle for its functional inactivation. Elife 6 (2017).

18. Preissler, S., Rato, C., Perera, L.A., Saudek, V. & Ron, D. FICD acts bifunctionally to AMPylate and de-AMPylate the endoplasmic reticulum chaperone BiP. Nat Struct Mol Biol 24, 23–29 (2017).

19. Das, D. et al. Crystal structure of the Fic (Filamentation induced by cAMP) family protein SO4266 (gi|24375750) from Shewanella oneidensis MR-1 at 1.6 A resolution. Proteins 75, 264–71 (2009).

20. Veyron, S., Peyroche, G. & Cherfils, J. FIC proteins: from bacteria to humans and back again. Pathog Dis 76 (2018).

21. Stanger, F.V. et al. Intrinsic regulation of FIC-domain AMP-transferases by oligomerization and automodification. Proc Natl Acad Sci U S A 113, E529–37 (2016).

22. Hegstad, K., Mikalsen, T., Coque, T.M., Werner, G. & Sundsfjord, A. Mobile genetic elements and their contribution to the emergence of antimicrobial resistant Enterococcus faecalis and Enterococcus faecium. Clin Microbiol Infect 16, 541–54 (2010).

23. Lebreton, F. et al. Tracing the Enterococci from Paleozoic Origins to the Hospital. Cell 169, 849–861 e13 (2017).

24. Casey, A.K. et al. Fic-mediated deAMPylation is not dependent on homodimerization and rescues toxic AMPylation in flies. J Biol Chem 292, 21193–21204 (2017).

25. Carafoli, E. & Krebs, J. Why Calcium? How Calcium Became the Best Communicator. J Biol Chem 291, 20849–20857 (2016).

26. Harms, A., Brodersen, D.E., Mitarai, N. & Gerdes, K. Toxins, Targets, and Triggers: An Overview of Toxin-Antitoxin Biology. Mol Cell 70, 768–784 (2018).

27. Bravo, R. et al. Endoplasmic reticulum and the unfolded protein response: dynamics and metabolic integration. Int Rev Cell Mol Biol 301, 215–90 (2013).

28. Krebs, J., Agellon, L.B. & Michalak, M. Ca(2+) homeostasis and endoplasmic reticulum (ER) stress: An integrated view of calcium signaling. Biochem Biophys Res Commun 460, 114–21 (2015).

29. Carreras-Sureda, A., Pihan, P. & Hetz, C. Calcium signaling at the endoplasmic reticulum: fine-tuning stress responses. Cell Calcium 70, 24–31 (2018).

30. Montero, M. et al. Monitoring dynamic changes in free Ca2+ concentration in the endoplasmic reticulum of intact cells. EMBO J 14, 5467–75 (1995).

31. Wong, W.L., Brostrom, M.A., Kuznetsov, G., Gmitter-Yellen, D. & Brostrom, C.O. Inhibition of protein synthesis and early protein processing by thapsigargin in cultured cells. Biochem J 289 (Pt 1), 71–9 (1993).

32. Romani, A.M. Cellular magnesium homeostasis. Arch Biochem Biophys 512, 1–23 (2011).

33. Xu, C., Bailly-Maitre, B. & Reed, J.C. Endoplasmic reticulum stress: cell life and death decisions. J Clin Invest 115, 2656–64 (2005).

34. Morris, G. et al. The Endoplasmic Reticulum Stress Response in Neuroprogressive Diseases: Emerging Pathophysiological Role and Translational Implications. Mol Neurobiol (2018).

35. Maly, D.J. & Papa, F.R. Druggable sensors of the unfolded protein response. Nat Chem Biol 10, 892–901 (2014).

36. Kabsch, W. Integration, scaling, space-group assignment and post-refinement. Acta Crystallogr D Biol Crystallogr 66, 133–44 (2010).

37. Vonrhein, C. et al. Data processing and analysis with the autoPROC toolbox. Acta Crystallogr D Biol Crystallogr 67, 293–302 (2011).

38. Adams, P.D. et al. PHENIX: a comprehensive Python-based system for macromolecular structure solution. Acta Crystallogr D Biol Crystallogr 66, 213–21 (2010).

39. Emsley, P. & Cowtan, K. Coot: model-building tools for molecular graphics. Acta crystallographica. Section D, Biological crystallography 60, 2126–32 (2004).

40. Morin, A. et al. Collaboration gets the most out of software. Elife 2, e01456 (2013).

